# Maternal exposure to cigarette smoking induces immediate and durable changes in placental DNA methylation affecting enhancer and imprinting control regions

**DOI:** 10.1101/852186

**Authors:** Sophie Rousseaux, Emie Seyve, Florent Chuffart, Ekaterina Bourova-Flin, Meriem Benmerad, Marie-Aline Charles, Anne Forhan, Barbara Heude, Valérie Siroux, Remy Slama, Jorg Tost, Daniel Vaiman, Saadi Khochbin, Johanna Lepeule, the EDEN mother-child cohort study group

**Affiliations:** Univ. Grenoble Alpes, Inserm, CNRS, IAB, 38000 Grenoble, France; Inserm U1153, Early Origins of Child Health and Development team, Research Center for Epidemiology and Biostatistics Sorbonne Paris Cité (CRESS), Paris Descartes University, Villejuif, France; Laboratory for Epigenetics and Environment, Centre National de Recherche en Gnomique Humaine, CEA – Institut de Biologie François Jacob, Evry, France; Genomics, Epigenetics and Physiopathology of Reproduction, Institut Cochin, U1016 Inserm – UMR 8104 CNRS – Paris-Descartes University, Paris, France

## Abstract

**Objective:** Exposure to cigarette smoking during pregnancy has been robustly associated with cord blood DNA methylation. However, little is known about such effects on the placenta; in particular, whether cigarette smoking before pregnancy could also induce epigenetic alterations in the placenta of former smokers is unknown.

**Design and results:** Placental DNA methylation levels were measured in 568 women and compared among non-smokers and women either smoking during their pregnancy or who had ceased smoking before pregnancy. An Epigenome Wide Association Study identified 344 Differentially Methylated Regions (DMRs) significantly associated with maternal smoking status. Among these 344 DMRs, 262 showed “reversible” alterations of DNA methylation, only present in the placenta of current smokers, whereas 44 were also found altered in former smokers, whose placenta had not been exposed directly to cigarette smoking. This observation was further supported by a significant demethylation of *LINE-1* sequences in the placentas of both current (−0.43 (−0.83 to −0.02)) and former smokers (−0.55 (−1.02 to −0.08)) compared to nonsmokers. A comparative analysis of the epigenome landscape based on the ENCODE placenta data demonstrated an enrichment of all 344 DMRs in enhancers histone marks. Additionally, smoking-associated DMRs were found near and/or overlapping with 13 imprinting gene clusters encompassing 18 imprinted genes.

**Conclusions:** DNA methylation patterns alterations were found in 344 genomic regions in the placenta of women smoking during their pregnancy, including 44 DMRs and *LINE-1* elements, where methylation changes persisted in former smokers, supporting the hypothesis of an “epigenetic memory” of exposure to cigarette smoking before pregnancy. Enhancers regions, including imprinting control regions were also particularly affected by placenta methylation changes associated to smoking, suggesting a biological basis for the sensitivity of these regions to tobacco exposure and mechanisms by which fetal development could be impacted.

## Introduction

Despite an increasing awareness of smoking associated risks in pregnancy and although smoking cessation is recognized as one of the most effective actions for improving mothers’ and children’s health (*1*), between 5 and 20 % of women continue to smoke during pregnancy in the United States and Europe (*2, 3*), with a prevalence of about 8% in Germany, 14% in Spain, 12% in UK and 17% in France (*3*). Maternal smoking during pregnancy is the most frequent preventable cause of adverse pregnancy outcomes (*4*) including placental abruption, placenta previa (*5*), preterm delivery (*6*) and some congenital anomalies (*7*). It has also been causally associated with intrauterine growth restriction (*8*). In the long term, maternal smoking in pregnancy is associated with adverse outcomes on child’s respiratory (*9, 10*) and cardiometabolic health (*11, 12*), neurodevelopment (*13*) and cancer (*14–16*). Despite this large amount of evidence supporting the effects of maternal smoking on the placenta, fetus and child, the molecular mechanisms involved in these effects remain poorly understood.

Intrauterine life is a critical period of plasticity during which environmental insults can alter the developmental programing via epigenetic phenomena, with immediate effects visible at birth or delayed effects that appear in childhood, puberty or adulthood (*17*). The most explored epigenetic mark so far has been the methylation of DNA, a modification known to be involved in the control of gene expression. More specifically, DNA methylation involves the addition of a methyl group to a cytosine, which in mammals is located upstream guanine residues (a “CpG” dinucleotide) on the DNA molecule. Modifications of DNA methylation can be replicated through cell divisions and can persist even in the absence of the cause that established them (biological memory) (*18*). Although DNA methylation is relatively stable in somatic cells, where it is transmitted through mitotic divisions, there are specific periods during development when the methylation pattern is reprogrammed, such as after fertilization in the pre-implantation embryo (*19*). Maternal smoking during pregnancy has been associated with DNA methylation levels in buccal cells of children offspring (*20*), in blood of adolescent offspring (*21*), and in peripheral blood granulocytes in adult women (*22*). In neonates, most studies on tobacco smoking during pregnancy have focused on cord blood DNA methylation (*23–26*). As a transient organ, the placenta is particularly interesting since it may provide a “molecular archive” of the prenatal environment (*27*), as recently shown for maternal smoking during pregnancy (*28*).

A few studies have investigated the association between placental DNA methylation and maternal smoking during pregnancy (*29–33*). Most studies were restricted to candidate CpG investigations or low-density genome-wide analyses and many of them were conducted on a small sample size. Importantly, these studies only included women who smoked throughout their pregnancy or who quitted smoking during the pregnancy. Smoking cessation is highly encouraged by doctors during pregnancy as well as prior to pregnancy. For both doctors and pregnant women, it is important to know whether exposure to tobacco smoking before and/or during pregnancy could differentially impact the offspring, and how. This work aimed at identifying cigarette-induced alterations in placental DNA methylation in women currently smoking during pregnancy and in women who quitted in anticipation of pregnancy. Placenta samples from 568 women were analyzed using Illumina HM450k BeadChip. A multidisciplinary approach combining statistical analyses with biological knowledge of the epigenome landscape enabled the identification and characterization of specific genomic regions differentially methylated in the placentas at birth as a result of cigarette smoking exposure during or prior to pregnancy (Figure 1).

**Figure 1:**
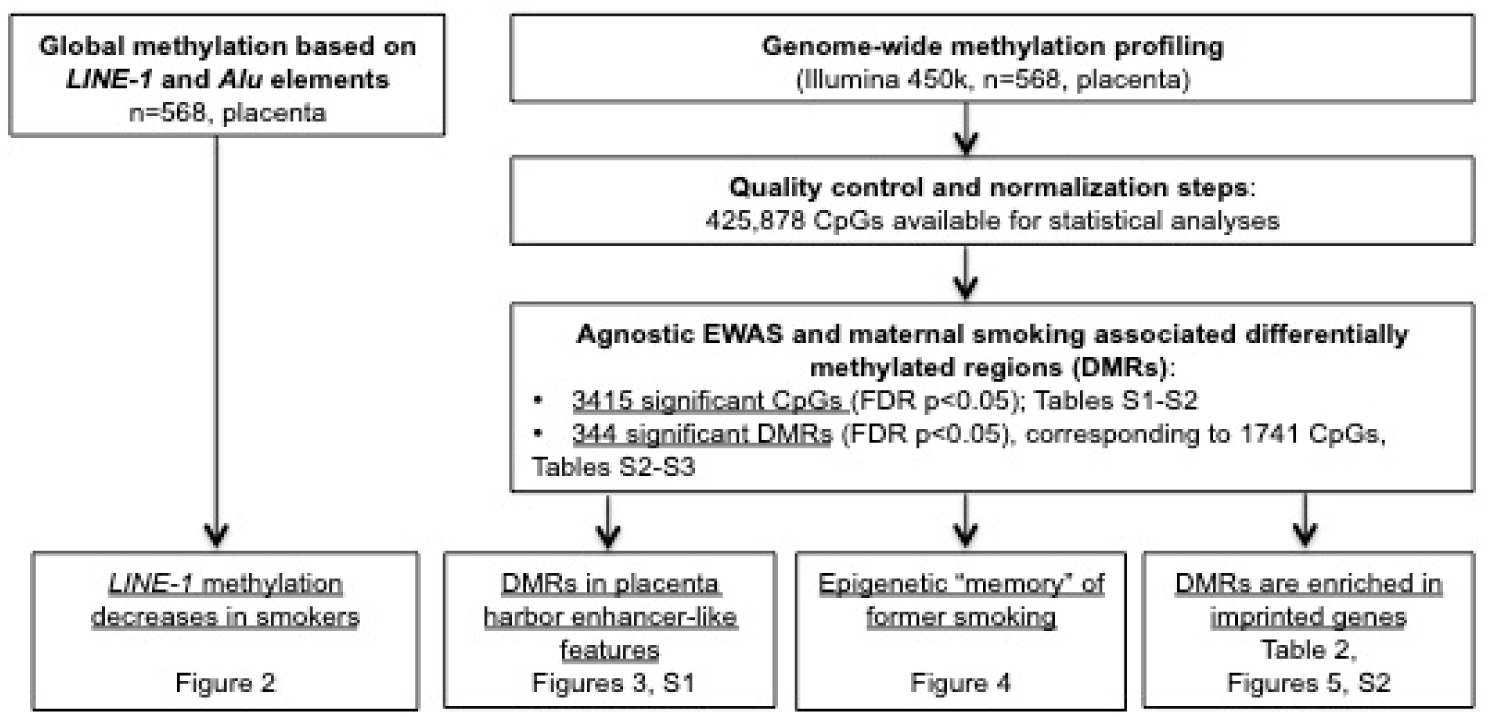
workflow of the study.

## Materials and Methods

### Study population

Study participants included in this analysis are a subset of the participants enrolled in the EDEN mother-child cohort between 2003 and 2006 (*34*). EDEN is a two-center study that has included 2002 pregnant women, mainly Caucasian, before 24 weeks of gestation in the university hospitals of Nancy and Poitiers, France. Exclusion criteria were multiple pregnancies, pre-pregnancy diabetes, French illiteracy and plans to move outside the region within the following 3 years. Lifestyle, demographic and medical data were collected by questionnaires and interviews during pregnancy and after delivery. DNA methylation (DNAm) was measured in placenta samples from 668 women (*35*). Placenta samples were collected at delivery by the midwife or the technician of the study using a standardized procedure. Samples of around 5mm x 5mm were excised from the middle of the placental fetal side and were immediately frozen at −80°C. The EDEN cohort received approval from the ethics committee (CCPPRB) of Kremlin Bicêtre and from the French data privacy institution “*Commission Nationale de l’Informatique et des Libertés”* (CNIL). Written consent was obtained from the mother for herself and for the offspring.

### Smoking exposure variables

Prenatal maternal cigarette smoking was collected by questionnaires administered by the midwives during prenatal and postpartum clinical examinations. *Nonsmoker* were defined as women who did not smoke during the 3 months before and during the pregnancy. *Current smokers* were defined as mothers smoking ≥1 cigarette per day throughout the duration of the pregnancy. All current smokers during pregnancy also smoked during the 3 months before pregnancy. *Former smokers* were defined as women who reported smoking during the 3 months preceding the pregnancy and declared not smoking throughout the duration of the pregnancy. Women who quitted smoking during pregnancy or cases with missing information regarding their smoking status at some point during the 3 months preceding the pregnancy or during the pregnancy were excluded, leaving 568 participants for this analysis.

### Placental DNA methylation measurements and quality control

DNA from placental samples was extracted using the QIAsymphony instrument (Qiagen, Germany). The DNA methylation analysis was performed by the Centre National de Recherche en Génomique Humaine (CNRGH, Evry, France). The DNA samples were plated onto 96-well or 48-well plates. In total, nine plates including 64 chips were used. These plates were analyzed in 4 batches. The ratios for sex (boy/girl) and recruitment centre (Poitiers/Nancy) were balanced for each chip. Fifteen samples were measured in quadruplicates and one sample in duplicate across batches, sample plates and chips to detect technical issues such as batch effects. The Illumina’s Infinium HumanMethylation450 BeadChip, representing over 485,000 individual CpG sites, was used to assess levels of methylation in placenta samples following the manufacturer’s instructions (Illuminas, San Diego, CA, USA). Raw signals of 450K BeadChips were extracted using the GenomeStudio^®^ software (v2011.1. Illumina). The DNA methylation level of each CpG was calculated as the ratio of the intensity of fluorescent signals of the methylated alleles over the sum of methylated and unmethylated alleles (β value). All samples passed initial quality control and had on average more than 98 % of valid data points (detection p-value <0.01). A refined version of the Subset Quantile Normalization (SQN) pipeline (*36*)including a revised annotation file (*37*) was used for data processing, correction and normalization. Data processing and normalization did not change the density distribution of the DNA methylation levels (results not shown). Intensity values were corrected for potential biases in fluorescent dye intensity and background corrected using the *lumi* R package (*38*) as implemented in the SQN pipeline. Probes potentially influenced by SNPs underlying the entire sequence of the probe (+1 or + 2 bases depending on the Infinium probe type) that are present in the EUR population of the 1000 Genome project (http://www.1000genomes.org)) at a frequency of more than 5% were removed from the analysis. Probes previously reported to map to several genomic regions were removed (*39*). The SQN pipeline uses the intensity signals of high-quality (i.e. low detection p-value) Infinium I probes as “anchors” to estimate a reference distribution of quantiles for probes in a biologically similar context based on the annotation file (*36*). This reference was then used to estimate a target distribution of quantiles for InfII probes as a means to provide an accurate normalization of InfI/InfII probes and correct for the shift. SQN is performed for each individual separately. A principal component analysis as well as a hierarchical clustering were applied and showed no overall difference in the methylation patterns across participants samples and replicates, so that a quantile normalization was performed for between sample normalization. After quality control and normalization steps, there were 426,049 CpG sites left. Methylation beta values ranged from 0 to 1. Data points with a detection p-value >0.01 were excluded from subsequent analyses. To reduce the influence of potential outliers, we excluded data points below the 25^th^ percentile minus 3 interquartile ranges or above the 75^th^ percentile plus 3 interquartile ranges for each probe, which removed 0.4% of all methylation beta values across participants. CpGs with more than 25% of missing data were removed, leaving 425,878 CpG sites for statistical analyses.

Global methylation was also evaluated by measuring methylation in four CpG sites of repetitive *Alu* elements (*Alu*) and long interspersed nucleotide elements 1 (*LINE-1*) using a previously published pyrosequencing methylation assay (*40*). We then used the median percent methylation of the four CpG sites.

### Cellular composition of placenta samples

Cellular composition of biological samples is a potential confounder in epigenetic epidemiological studies. In the absence of reference methylomes for placental tissue, we used a reference-free method, the RefFreeEWAS package available in R (*41*), to deconvolute cell-type proportions from DNA methylation array data. The method relies on the identification of latent variables as surrogates for cell-type mixture. From the 10,000 most variable CpGs, we identified the optimal number of cell-types to be 6. We then used the 425,878 CpGs to estimate the proportion of each cell-type per sample.

### Analytical approach

We hypothesized that maternal cigarette smoking during or before pregnancy could alter the placental function through modifications of DNA methylation. Therefore, we investigated the relationship of maternal cigarette smoking with global DNA methylation and gene-specific methylation (Figure 1). In order to identify potentially relevant changes in genomic methylation sites, we performed an Epigenome-Wide Association Study (EWAS) and identified differentially methylated regions (DMR). We then focus our subsequent analyses on these smoking associated DMRs and corresponding CpGs. We first characterized the epigenetic context of the regions whose DNA methylation patterns were significantly affected by cigarette smoking. Second, we classified the DMRs and CpGs into two categories labeled as “reversible” and “memory”. Finally, we looked for proximity and/or overlaps between the DMRs and the known imprinting control regions and promoter regions in order to assess the potential impact of tobacco exposure related methylation alterations of our DMRs on the regulated expression of these genes (Figure 1).

### Statistical analysis for the EWAS and identification of DMR

We studied the association between smoking status (nonsmoker, current, former) and both the repetitive elements *Alu* and *LINE-1* and the CpG-specific methylation level using the following robust linear regression model in order to account for potential outliers and heteroscedasticity:

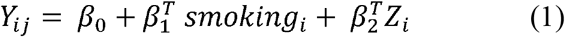

where Y_ij_ is the methylation measurement for CpG or repetitive element *j* in subject *i*, smoking_i_ is the smoking status of subject i, Zi is the set of *a priori* selected adjustment factors including: child sex, parity (0, 1, ≥2 children), maternal age at end of education (≤18, 19-20, 21-22, 23-24, ≥25 years), season of conception, study center (Poitiers and Nancy), maternal body mass index (BMI) before pregnancy (≤18.5, 18.5-25, 25-30, ≥30), maternal age at delivery (linear and quadratic terms), gestational duration (linear and quadratic terms), technical factors related to the methylation measurements (batch, plate and chip) and estimated cell-type proportions. We applied the Benjamini and Hochberg False Discovery Rate (FDR) correction to the p-values to account for multiple testing (*42*). The FDR corrected p-values were calculated for the 425,878 CpGs for the agnostic EWAS. An FDR-corrected p-value <0.05 was considered statistically significant.

To identify Differentially Methylated Regions (DMRs) from the EWAS (425,878 CpGs), we used comb-p, a method relying on the Stouffer-Liptak-Kechris correction that combines specific CpG p-values using sliding windows and accounting for correlation between CpGs (*43*). DMR p-values were adjusted for multiple testing by Šidák correction (*43*). Significant DMRs (p-value < 0.05) included at least 2 probes (p-value < 0.001) within a window of 2000 bp.

We compared our results from the EWAS and regional analysis with previous findings from studies investigating the relationship between maternal smoking in pregnancy and placenta and cord blood DNA methylation changes with the Illumina HM450k BeadChip.

All analyses were performed using the statistical software R (version 3.0.1) (R Core Team, 2013) and Python (version 2.7.14).

### Characterization of epigenetic context of smoking-associated DMRs

We investigated whether the smoking associated DMRs could be associated with specific features in their global epigenomic landscape. Indeed, the identification of a specific epigenome environment could not only explain the particular sensitivity of these regions but also help us anticipating the biological consequences of methylation alterations on the epigenomic control of placenta development. To address this question, we first identified a set of random regions of similar sizes to compare to our smoking associated DMRs. To identify “random DMRs”, smoking status was randomly assigned to each participant of our study in an iterative process repeated 122 times. The EWAS and DMRs analyses were performed for each dataset provided by each of the 122 iterations. This process identified 890 differentially methylated regions associated with smoking status just by chance.

We then looked for the epigenetic marks normally associated with these regions in placenta by using the corresponding ENCODE ChIPseq data (https://www.encodeproject.org, (*44*)). We compared the enrichment of our smoking associated DMRs to that of the 890 random DMRs for the following epigenetic marks: the post-translational modifications (PTM) of histone H3, mono and trimethyl lysine 4 (H3K4me1 and me3) and the acetylation of lysine 27 (H3K27ac). The files described in supplemental table S2 corresponding to the read counts for the presented histone H3 PTMs were retrieved from ENCODE (www.encodeproject.org) and used to produce heatmaps and profiles to map these modifications on our smoking associated DMRs and random DMRs. The deepTools softwares (deeptools.readthedocs.io) were used with options *computeMatrix reference-point --referencePoint TSS --binSize 50 --beforeRegionStartLength 2000 -- afterRegionStartLength 2000 --sortRegions keep* to extract PTM profiles.

### Identification of reversible and epigenetic memory methylation profiles among CpGs and DMRs

Smoking associated DMRs and corresponding CpGs were classified into categories, defined by the variations of the methylation patterns between the three groups of women, current smokers, nonsmokers and former smokers. First, by considering the regression coefficients and p-values of the three smoking groups for each of differentially methylated CpG, we labeled each CpG according to its pattern of association with smoking status as “reversible” (alteration present in current smokers but absent in the other two groups) or “memory” (alteration present in current smokers as well as in former smokers, compared to nonsmokers) (Figure 4 A and C). Second, each of the 344 DMRs was assigned to the category most represented among its CpGs labels and corresponding to at least 50% of its CpGs (Figure 4 B and C). The other DMRs, including those containing the same proportions of CpGs with different labels, were labeled “undefined”.

### Identifying imprinting control regions potentially affected by exposure to tobacco

We then addressed the question of whether the alterations of the DNA methylation patterns induced by cigarette smoking could have consequences on the regulation of the expression of imprinted genes. As opposed to most genes, which show bi-allelic expression (from both paternal and maternal alleles), imprinted genes are expressed only on one of the two parental alleles. This monoallelic expression is known to be controlled by the allelic differential DNA methylation on regions nearby or more distant, known as imprinting control regions (ICR). In the human, approximately 310 imprinted genes have been identified so far, which tend to group in clusters with shared ICR. Presently, 246 and 166 imprinted genes are respectively recorded in www.geneimprint.com and igc.otago.ac.nz databases, with 102 genes common to the two databases. In order to assess the potential impact of methylation alterations in our smoking associated DMRs on the regulated expression of these genes, we looked for proximity and/or overlaps between our smoking associated DMRs and the known ICRs and promoter regions associated with these genes. The candidate ICRs were defined by systematically matching the genes of the Illumina HM450k BeadChip annotations corresponding to smoking associated DMRs and the gene annotations present on databases *MetaImprint (*http://202.97.205.76:8080/MetaImprint/, (*45*)*), geneimprint (*http://www.geneimprint.com, (*46*)*) and igc.otago (*http://igc.otago.ac.nz, (*47))*.

## Results

### Population characteristics

On average (±SD), the participating mothers were 29.3 (± 5.0) years old, with a pre-pregnancy body mass index of 23 (± 4.2) kg/m² (Table 1). Mean gestational duration was 39.7 (±1.7) weeks and 30 babies (5.3 %) were born preterm (<37 gestational weeks). Among the 568 women participating to this analysis, 381 (67.1 %) were nonsmokers (i.e. did not smoke in the 3 months before pregnancy nor during the pregnancy), 117 (20.6 %) were current smokers (i.e. did smoke ≥1 cigarette per day throughout the duration of the pregnancy), and 70 (12.3 %) were former smokers (i.e. did smoke in the 3 months preceding the pregnancy and stopped before the pregnancy).

**Table 1.** Characteristics of the EDEN study population (n=568)

| Characteristics | mean±SE | n (%) |
| --- | --- | --- |
| **Center** |  |  |
| Poitiers |  | 231 (41) |
| Nancy |  | 337 (59) |
| **Sex of offspring** |  |  |
| Male |  | 293 (52) |
| Female |  | 275 (48) |
| **Parity** |  |  |
| 0 |  | 239 (42) |
| 1 |  | 226 (40) |
| ≥2 |  | 102 (18) |
| Missing |  | 1 (0) |
| **Maternal age at end of education (year)** |  |  |
| ≤18 |  | 105 (18) |
| 19-20 |  | 89 (16) |
| 21-22 |  | 127 (22) |
| 23-24 |  | 134 (24) |
| ≥25 |  | 113 (20) |
| **Season of conception** |  |  |
| January – March |  | 120 (21) |
| April – June |  | 129 (23) |
| July – September |  | 157 (28) |
| October – December |  | 162 (29) |
| **Pre-pregnancy Body Mass Index (kg/m<sup>2</sup>)** |  |  |
| <18.5 |  | 60 (9) |
| 18.5-25 |  | 442 (66) |
| 25-30 |  | 112 (17) |
| ≥30 |  | 40 (6) |
| Missing |  | 14 (2) |
| **Maternal age (year)** | 29.3±5.0 |  |
| **Gestational duration (weeks)** | 39.7±1.7 |  |
| **Maternal smoking<sup>a</sup>** | 1.8±5.0 |  |
| Nonsmoker |  | 381 (67) |
| Current smoker |  | 117 (21) |
| Former smoker |  | 70 (12) |

### Maternal cigarette smoking was associated with lower LINE-1 methylation levels

The average methylation level was 26.1 (±1.9) for *LINE-1* and 16.1 (±1.0) for *Alu*. Women who smoked had significantly lower methylation levels for *LINE-1* compared to non-smoking women, with a slightly larger association for former smokers (−0.55 (−1.02; −0.08)) than for current smokers (−0.43 (−0.83; −0.02)) (Figure 2). This effect was not observed for *Alu* elements.

**Figure 2:**
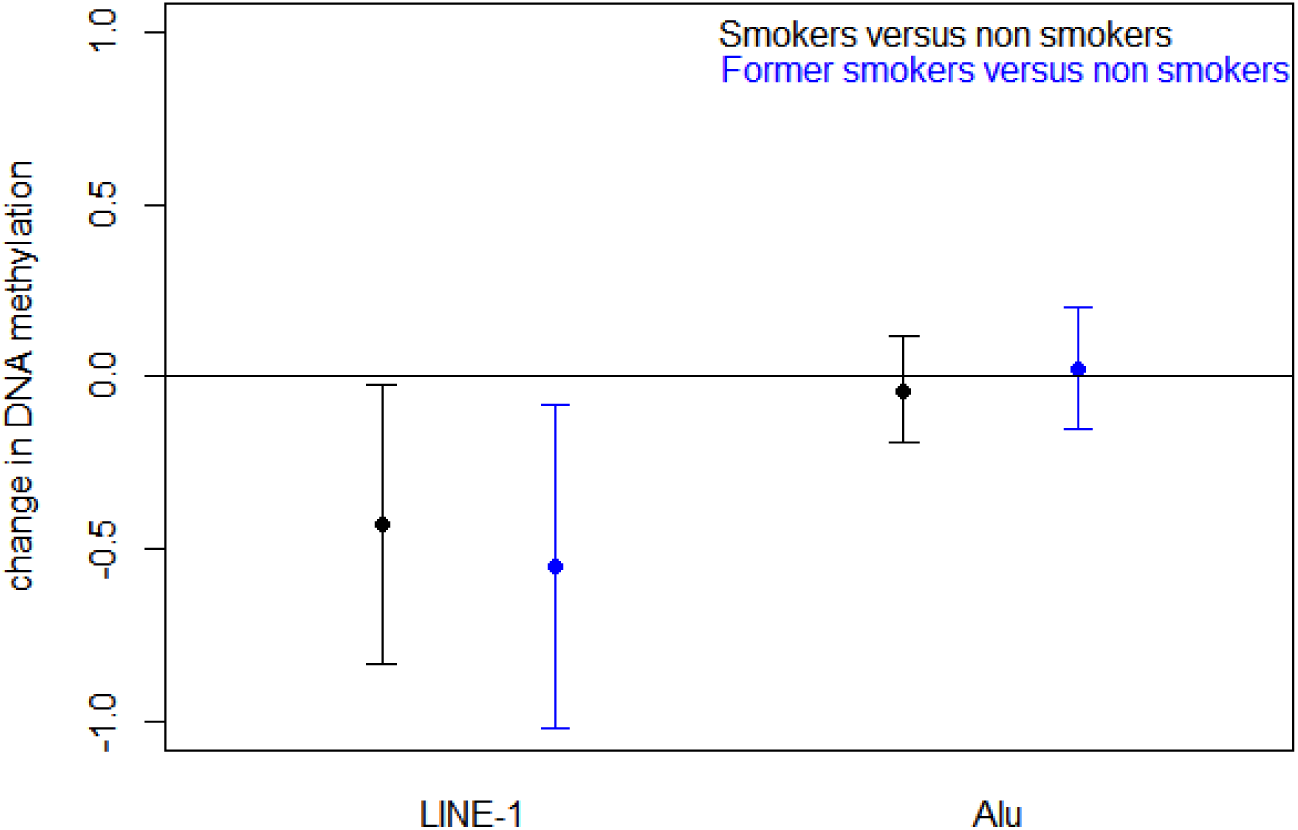
Adjusted association between smoking status and repetitive DNA methylation elements.

### 344 genomic regions were differentially methylated according to maternal smoking

Among the 425,878 CpGs explored in the adjusted Epigenome Wide Analysis Study (EWAS) (Supplemental table S1), 344 Differentially Methylated Regions (DMRs) were identified (Supplemental table S2). These 344 DMR included a total of 1741 CpGs (471 of which were individually significant (FDR p-value <0.05) in the EWAS). We focused our subsequent analyses on these 344 DMRs and their associated 1,741 CpGs.

### Smoking-associated DMRs were depleted in gene promoters and harbor enhancer-like features

The analysis of ENCODE placental data comparing specific epigenetic marks between the 344 smoking associated DMRs and randomly selected regions revealed a relative depletion in H3K4me3, which is considered a hallmark of gene promoter regions, as well as a relative enrichment in the H3K4me1 and H3K27ac in our smoking related DMRs (Figure 3).

**Figure 3:**
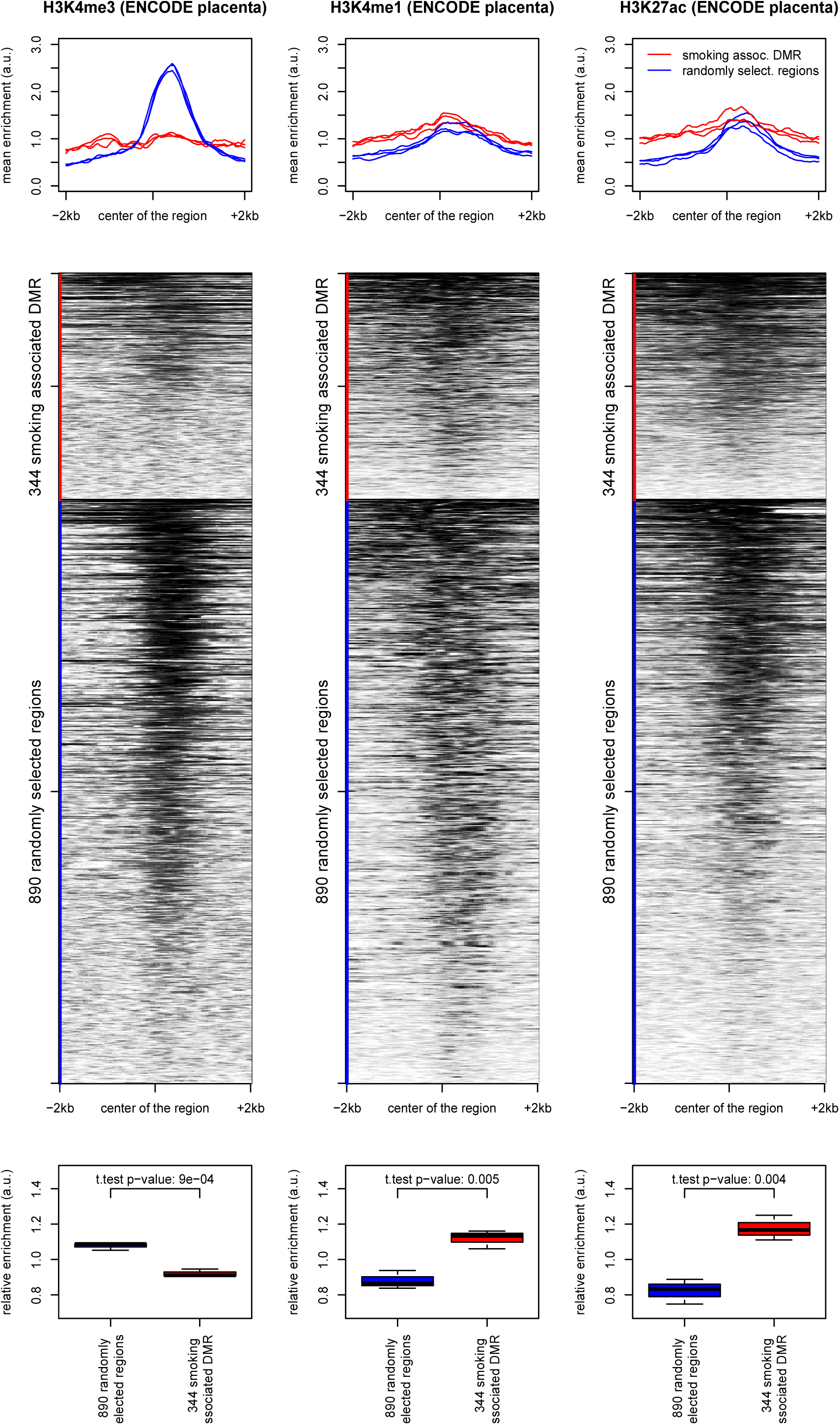
Analysis of the placenta ChipSeq ENCODE data. The 344 DMR associated with pregnancy smoking status are depleted in H3K4me3 and enriched in H3K4me1 and H3K27ac, compared to 890 randomly selected regions of similar sizes on the Illumina array. The curves (A) show the mean profiles of ChipSeq read counts in our 344 DMRs (red) and in the 890 random regions (blue) in triplicates. The heatmaps (B) correspond to the ChipSeq data of one representative placenta sample for each mark. The box plots (C) show the distribution of areas under the corresponding curves. Heatmaps corresponding to all three samples are shown in supplemental files (Supplementary Figure S1).

### 44 DMRs bear “epigenetic memory” of maternal smoking before pregnancy

A total of 262 DMRs were bearing “reversible” alterations, only found in the current smokers group, whereas 44 DMRs showed alterations of placental DNA methylation not only in current smokers but also in former smokers whose placenta had not been exposed directly to cigarette smoking. The latter DMRs were therefore labeled as “epigenetic memory”.

### DMRs sensitive to smoking exposure were enriched in imprinted genes

The 344 smoking-associated DMRs were found near and/or overlapping with 13 imprinting control regions (ICR) encompassing 18 imprinted genes. An analysis of the expression patterns of these genes in normal tissues (Table 2, supplemental Figure S2) shows that most of these genes have a relatively high expression in placenta, supporting their important role during placenta development. Three of the 344 DMRs whose methylation levels were altered by exposure to smoking were found consistently close to (<1kb) imprinted loci (known ICR), regardless of the study used as a reference for ICR localization (Figure 5). The first two loci, NNAT/BLCAP (20q11.23) and SGCE/PEG10 (7q21.3) were both found associated with a reversible alteration of DNA methylation, which was respectively decreased or increased for all CpGs (but 2 for NNAT/BLCAP) in women currently smoking during pregnancy, compared to both former and nonsmokers. These two DMRs were the longest ones identified, with 33 and 37 CpGs respectively for NNAT/BLCAP and SGCE/PEG10. Interestingly, the third locus, H19/MIR675 (11p15.5), was affected by a decreased methylation not only in placenta of currently smoking pregnant women, but some methylation alteration was also detectable in placenta that had never been directly exposed to cigarette smoking.

**Table 2:** too large for being included in the manuscript. Please see excel file named table 2.

**Figure 4:**
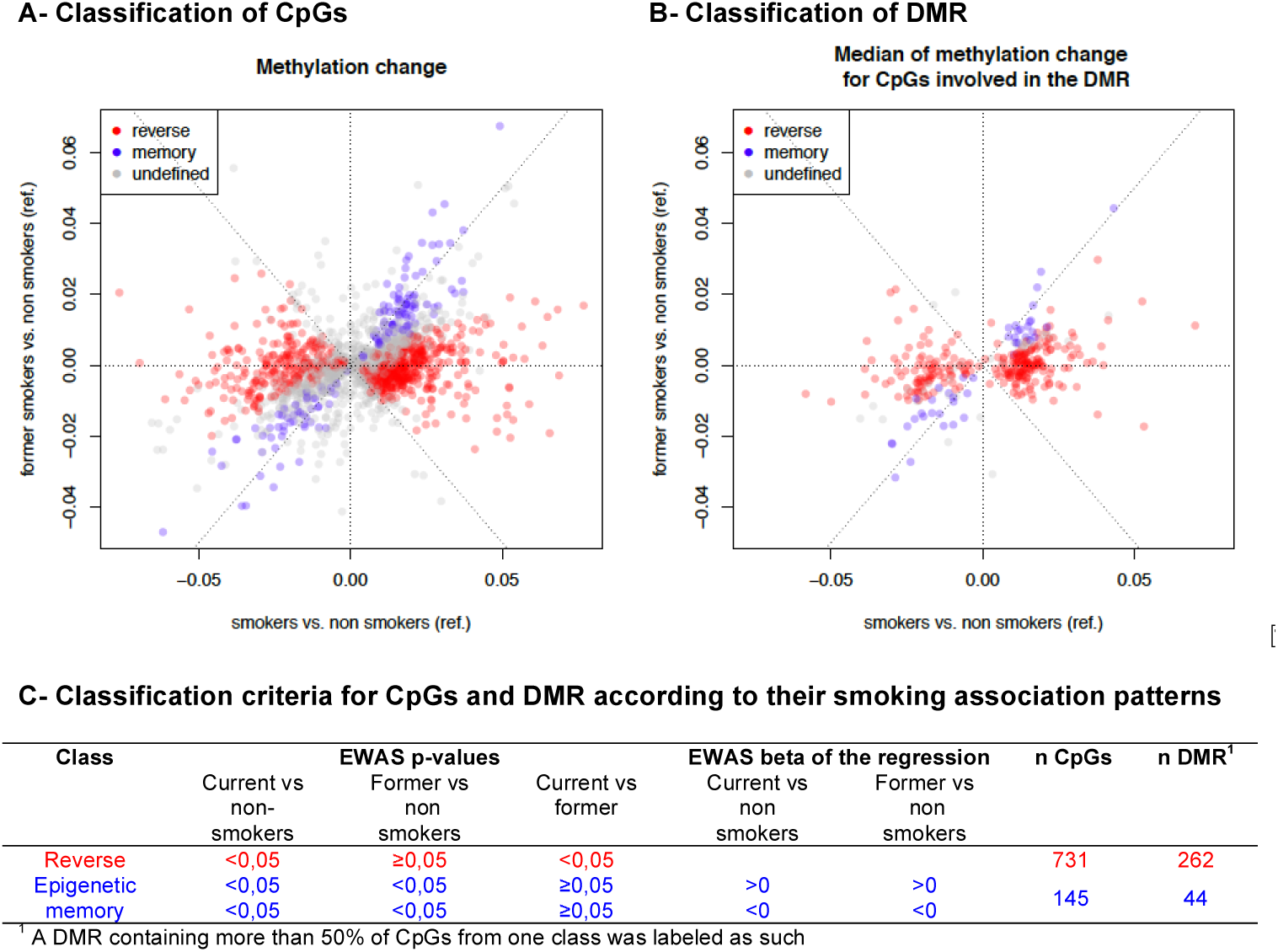
Classification of “epigenetic memory”, “reversible” CpGs and DMR according to their smoking association methylation patterns.

**Figure 5:**
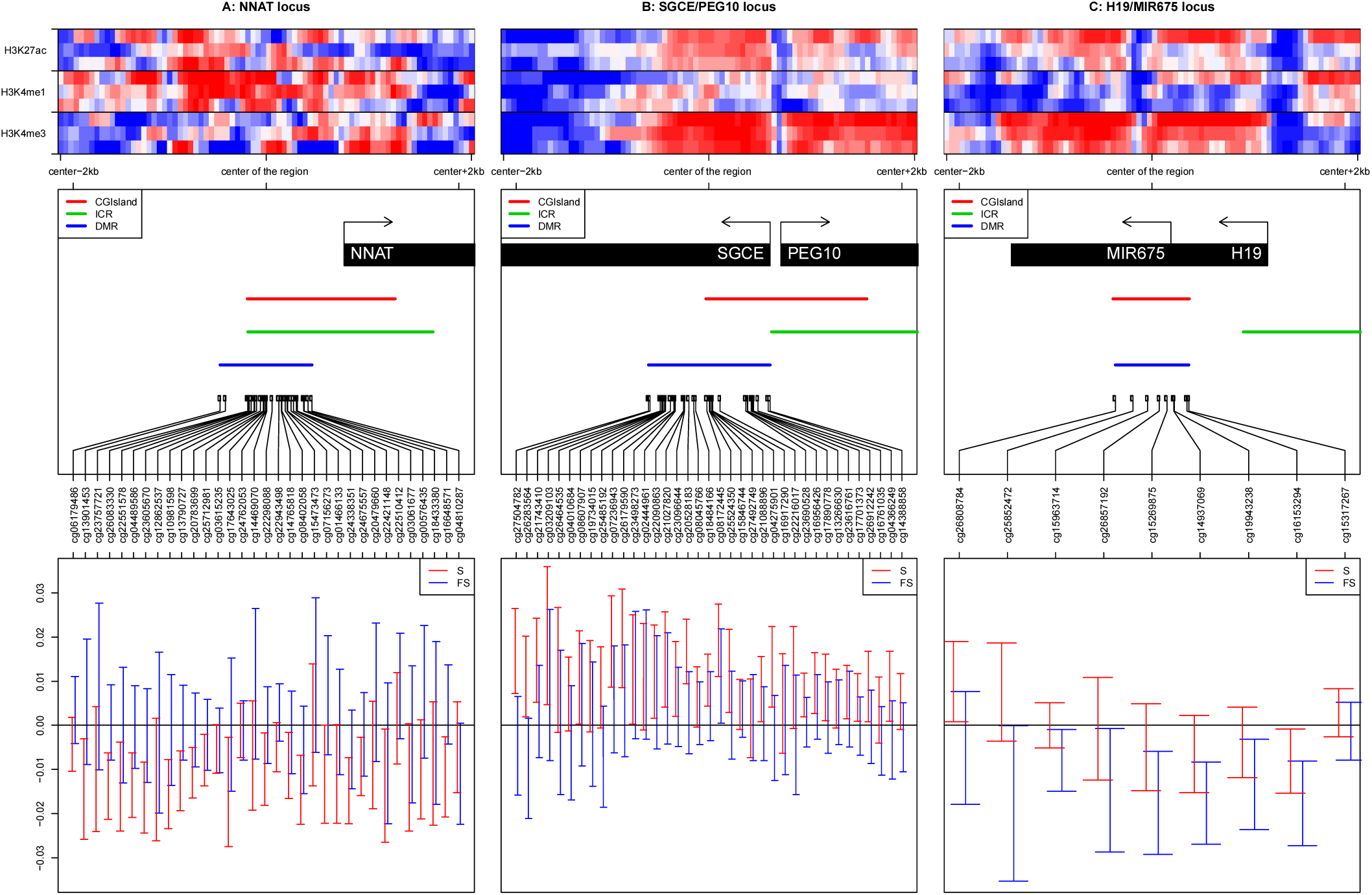
The methylation levels of three imprinted loci are consistently modified following exposure to cigarette smoking. The left, center and right panels are respectively centered on the DMRs associated with the NNAT locus +/- 2kb, the SGCE/PEG10 locus +/- 2kb and the H19/MIR675 locus +/- 2kb. The top panels show heatmaps respectively corresponding to H3K4me3, H3K4me1 and H3k27ac enrichments in placenta around the center of the region of interest +/- 2 kilobases. For each mark we downloaded triplicates from ENCODE data (see methods). We computed the enrichement matrices from bigWig files using the deepTools software and displayed it using a custom R script. The middle panels show genes and CGislands from the Illumina Infinium Human Methylation 450K BeadChip annotations, ICR from Pervjakova et al. (*73*), the DMR we identified and thecorresponding Illumina probes of the region of interest. The bottom panels show the methylation changes in smokers (S) (resp. Former Smokers (FS)) in red (resp. blue) compared to nonsmokers. The points represent the beta of the linear model and the error bars correspond to more or less 2 standard deviations.

## Discussion

The present study identified 344 genomic regions that were significantly differentially methylated in the placenta according to cigarette smoking during pregnancy. Interestingly, in 44 of these DMRs, we observed persistent methylation changes in the placentas of former smokers, despite an absence of direct exposure of these placentas to tobacco, suggesting the possibility of an “epigenetic memory” of exposure to cigarette smoking before pregnancy. This result is also supported by the observation of a significant demethylation of *LINE-1* sequences in the placentas of former smokers. Exploration of the epigenetic status of all DMRs revealed that genomic regions bearing “enhancer-like” epigenetic marks, namely histone post-translational modifications H3K4me1 and H3K27ac, are enriched among our 344 DMRs, suggesting that placenta enhancer genomic regions could be particularly sensitive to tobacco exposure. Furthermore, tobacco-associated DMRs also overlap with genomic regions controlling imprinted genes, known to have an important role in fetal and placental development.

### Strengths and limitations

Potential limitations of our study relate to the characterization of the smoking phenotype. Indeed, maternal smoking was evaluated by questionnaires administered by midwives involved in the study, which might lead to an underestimation of the number of smoking women and the effect of smoking, due to under-reporting of tobacco consumption. However, on a subsample of 100 women whose cotinine levels were measured in urine between 24 and 28 gestational weeks (*50*), we found only one former-smoker with cotinine levels potentially compatible with current smoking (> 50 ng/mL) suggesting that self-reporting was a reasonably accurate indicator of the maternal smoking status in our cohort. Another limitation inherent to this study concerns the relatively small size of the former smokers group, which could result in a lack of statistical power and affect the significance of CpG methylation differences between current and former smokers. Such case could lead to a misclassification of some genomic regions altered by cigarette exposure that could be categorized in the “epigenetic memory” group and rather belongs to the “reversible” group. Finally, the former smokers group likely includes women who stopped smoking in anticipation of a pregnancy (i.e. before the conception) and women who stopped smoking in early pregnancy once they knew they were pregnant (i.e. a few days / weeks after conception). Time since quitting smoking was not available in our data. However, this might influence the methylation levels and would be an interesting piece of information to add to future research to provide insights into the existing risk faced by former smokers even months or years after cessation.

The present study includes the largest sample size of placentas investigated to date, which were collected in the context of a longitudinal cohort with repeated data on maternal smoking before and during pregnancy. The placenta is considered as an accurate ‘record’ of children’s in-utero exposures (*48*) and represents only a partial barrier, since many chemicals, such as polycyclic aromatic hydrocarbons, can pass through and reach the fetus(*49*). Additionally, this study is the first observation and characterization of DNA methylation alterations in the placentas of former smokers.

### Comparison with other studies

Only one study previously investigated placenta DNA methylation in relation to maternal smoking using a high density genome-wide approach (*33*). Out of the 1741 CpGs included in our 344 regions sensitive to smoking, 10 CpGs were also found associated (FDR corrected p-value <0.05) with maternal smoking by Morales et al (Supplemental table S2 and S3). These 10 CpGs are located on the following genes: *TRIO, CMIP/PLCG2, TINAGL1, SLC45A4/LINC01300, PDXK, ACOX3, TGM1, PDGFB/RPL3*, and are all included in DMRs associated with a reversible methylation pattern upon smoking cessation. *TRIO* (Trio Rho Guanine Nucleotide Exchange Factor) was reported to interact with benzo(a)pyrenes, resulting in decreased gene expression (*33*), which could be in accordance with the increased methylation levels observed in relation to maternal smoking exposure in both, Morales et al. and our study. *TINAGL1* (tubulointerstitial nephritis antigen like 1) and *PDGFB* (platelet derived growth factor subunit B) are broadly expressed in the placenta and have been identified as pro-angiogenic factors (*51*). Angiogenesis is a major process in pregnancy and *PDGFB* is likely to play an important role in the maintenance of utero-placental homeostasis (*52*). Nicotine, one of the thousands of compounds of tobacco smoke, is also a pro-angiogenic factor (*53*). In our results, *TINAGL1* was hypomethylated and *PDGFB* was hypermethylated in smokers compared to nonsmokers. In a recent study, *TINAGL1* was found downregulated in the placenta of pre-eclamptic pregnant women compared to normotensive women (*54*). Although further studies are required in order to determine the functional relationship between these genes and their placental methylation, our findings might be of interest in the search for explanations of the apparent protective effect of maternal smoking on the risk of pre-eclampsia, and more generally in the relationship between maternal tobacco smoking and alterations in placental angiogenesis (*55*).

Several studies identified CpGs altered in cord blood samples following exposure to tobacco smoking during pregnancy (*23–25*). Although some of our CpGs in placenta overlapped with the results of these studies, the direction for the associations was somewhat inconsistent (Supplemental table S2 and S3). This lack of agreement between placenta and cord blood can be explained by the fact that placenta and cord blood cells have different epigenetic signatures reflecting their different functions (*56*). However, a few CpGs were hypermethylated in cord blood as well as in our placentas of smoking women, including cg05549655 (*23–25*) cg00213123, cg23727072, and cg26516004 (*24*) located on *CYP1A1*.

### Possible explanations

An interesting observation is the demethylating effect of exposure to tobacco smoking on LINE-1 sequences. Compared to other tissues, the human placenta is known to have lower levels of *LINE-1* and *Alu* methylation levels (*57, 58*). In our study, these dispersed repetitive sequences were differentially affected since only *LINE-1* sequences showed decreased methylation levels as a consequence of exposure to tobacco whereas the methylation levels of *Alu* sequences were not significantly affected. Although we do not have yet an explanation for this difference, this observation suggests that different regions of the genome could have differential sensitivities to environmental cues. Another startling observation at this stage was that the *LINE-1* sequences were also significantly demethylated in the placenta of women not currently smoking but who were smoking before their pregnancy, suggesting that the epigenetic profiles of these placentas somehow could bear the “memory” of past-exposure to tobacco. Only a few studies had so far investigated global methylation in the placenta. One study of approximately 40 first trimester placentas, did not show any effect of maternal smoking on neither *AluYb8* nor *LINE-1* methylation levels (*59*). In another study of 379 term placentas, *AluYb8* methylation level was significantly higher in smoking women compared to nonsmokers, while no evidence of significant association was found for *LINE-1* (*60*). Another study conducted on 96 term placentas failed to find any association between *LINE-1* methylation and maternal smoking (*61*). These apparent discrepancies with our results could be explained by a lack of statistical power to detect an effect of smoking for the first study and for the other two the fact that the samples were collected preferentially on the maternal side of the placenta, as opposed to our study conducted on samples representing the fetal side of the placenta. Our observation of an effect of tobacco exposure on placental *LINE-1* methylation levels, either directly or via the establishment and transmission of a memory, was confirmed and extended by the high-resolution genome-wide methylome analysis with the Illumina 450K array.

Although a few studies have investigated the effect of time since quitting smoking on blood DNA methylation (*62, 63*), the potential effect of past smoking on the methylation of the placenta had not been explored. We focused on women who had smoked but quitted smoking before their pregnancy and therefore before the differentiation of the placenta. Our results identified 262 DMRs where the DNA methylation profiles were altered in the placenta of women currently smoking during their pregnancy and not in the placenta of nonsmokers or former smokers, suggesting that the DNA methylation alterations of these regions could be associated with direct exposure of the placenta to tobacco, and “reversible” upon smoking cessation. Another 44 regions showed altered DNA methylation profiles in the placenta not only in women actively smoking during their pregnancy but also in women exposed to cigarette smoking priori to their pregnancy. This result, in line with the significant demethylation of *LINE-1* sequences observed in the placenta of current and former smokers, suggests that the placentas of former smokers bears epigenetic marks reflecting a past exposure to smoking, prior to the development of the placenta. It implies that somehow these epigenetic marks were transmitted to the maternal cells from which the placenta originated and remained stable all through placenta differentiation. Beyond this particular context, this result suggests that the information of a past event could be durably printed in our epigenome, enabling the memory of the event to be carried through generations of cells all through their differentiation. This observation raises new fundamental questions in the field of epigenetics such as the mechanisms and molecular basis involved in such a transmission, which are presently totally unknown.

Since DNA methylation is only one of the many chemical marks involved in shaping the epigenome, we wished to explore other elements of the epigenetic landscape of the DMRs identified as associated with maternal smoking status. The question was whether some other elements of the epigenetic context of these DMRs could be associated with the particular sensitivity of these regions to smoking exposure. A meta-analysis of ENCODE data from placenta enabled us to explore the epigenome landscape normally associated with the most affected genomic regions. Interestingly, we found that most regions with smoking induced altered DNA methylation profiles are depleted in histone post-translational modifications (PTM) marks normally associated with poised genes promoter regions, such as H3K4me3, but are enriched in PTM generally associated with “enhancers” regions. This observation suggests that this epigenetic context is somehow related to a higher sensitivity of these regions to tobacco exposure.

Another major observation from this work is that tobacco induced DNA methylation alterations also affect imprinting control regions (ICR), whose allelic differential methylation normally controls the allele-specific expression of clusters of “imprinted” genes. A recent work highlighted the incomplete erasure of germline DNA methylation in human placenta and the effect of some allele-specific DNA methylation pattern on the expression of imprinted genes in placenta (*64*). Interestingly, three of our DMRs regions whose methylation profiles were altered by cigarette smoking exposure were found close to (<1kb) three imprinting loci, whose ICR are well defined. This finding suggests that the imprinted genes controlled by these ICR could have their expression patterns directly affected by smoking, and potential mechanisms by which tobacco could impact the epigenome and fetal growth. The first two loci, *NNAT/BLCAP* (20q11.23) and *SGCE/PEG10* (7q21.3) were both associated with altered (decreased and increased, respectively) DNA methylation in women currently smoking during pregnancy and classified as “reversible” regions. Increased expression of *NNAT/BLCAP* was associated with increased risk of large or small for gestational age infants (*65*). As for *SGCE/PEG10*, its methylation in cord blood was associated with birth weight (*66, 67*) and its expression in bronchial epithelium was associated with tobacco smoking in adults (*68*). The demethylation of the third locus, which controls the expression of *H19/MIR675* (11p15.5), was observed not only in the placenta of currently smoking women but also detectable in the placenta of former smokers suggesting that this important imprinted locus could be part of those bearing the memory of past-exposure to tobacco. Hypomethylation of this gene has been found associated with increased expression in fetal growth-restricted placentas (*69*). In a recent study, hypomethylation of *H19* in cord blood was found significantly associated with maternal smoking during pregnancy (*70*). Furthermore, placental expression of *H19* was associated with large for gestational age infants (*65*) and genetic variants have been identified and related to birth weight (*71, 72*). Our finding of a demethylation of the *H19/MIR675* locus in the placenta exposed to cigarette smoking, combined with previous observations, strongly support the hypothesis that the *H19/MIR675* locus is a major determinant of fetal growth and that cord blood and placental DNA methylation are both sensitive to direct smoking exposure.

## Conclusion and implications

This study investigating the effect of smoking on human placental DNA methylation at high resolution in the largest sample size published to date, has led to the identification of differentially methylated regions (DMRs) in the placentas of current smokers as well as former smokers. These tobacco-induced DMRs are enriched in epigenetic marks corresponding to enhancer regions, and some of them overlap regions controlling the monoallelic expression of imprinted genes, suggesting mechanisms by which tobacco could impact the epigenome and affect placental development and fetal growth. Additionally, altered DNA methylation patterns were not only observed in the placenta directly exposed to cigarette-smoking, but some alterations of DNA methylation patterns were also observed in the placenta of women who had smoked but quitted smoking in anticipation to pregnancy, suggesting the establishment of a “memory” of exposure to tobacco and transmission of epigenetic marks to placentas that had never been directly exposed to smoking. Hence this work brings new concepts and questions in the field of human epigenetics and suggests mechanisms by which an exposure to environmental cues could have not only direct but also long-term effects on human health. In addition to the important potential impact on our current knowledge of epigenetic transmission, our results bring essential information in terms of public health concerning potential long-term detrimental effects of smoking in young women, which could affect their offspring even after smoking cessation. This scientific report should support primary and secondary prevention of smoking associated health effects.

## Supplementary Materials

**Supplementary Figure S1:**
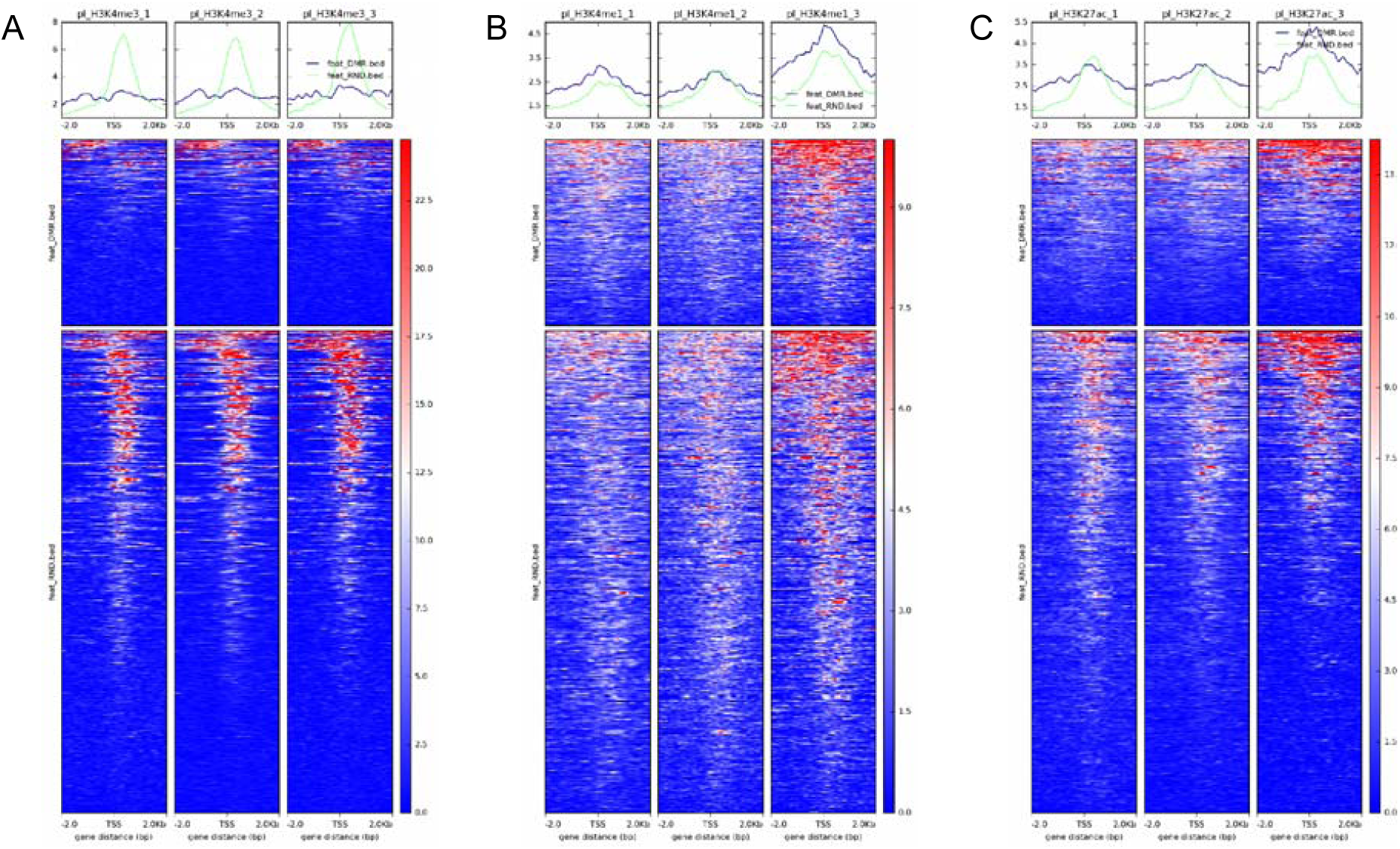
H3K4me3 (A), H3K4me1 (B) and H3K27ac (C) ChIPSeq signal of 3 placenta replicates. The top panels represent the mean signal values centered on our 344 DMRs +/-2kb (blue) and the mean signal values centered on the 890 random regions +/-2kb (green). The center panels represent heatmaps of the corresponding ChipSeq signals centered on our 344 DMRs +/-2kb (upper heatmaps) or on the 890 random regions+/-2kb (lower heatmaps).

**Supplementary Figure S2:**
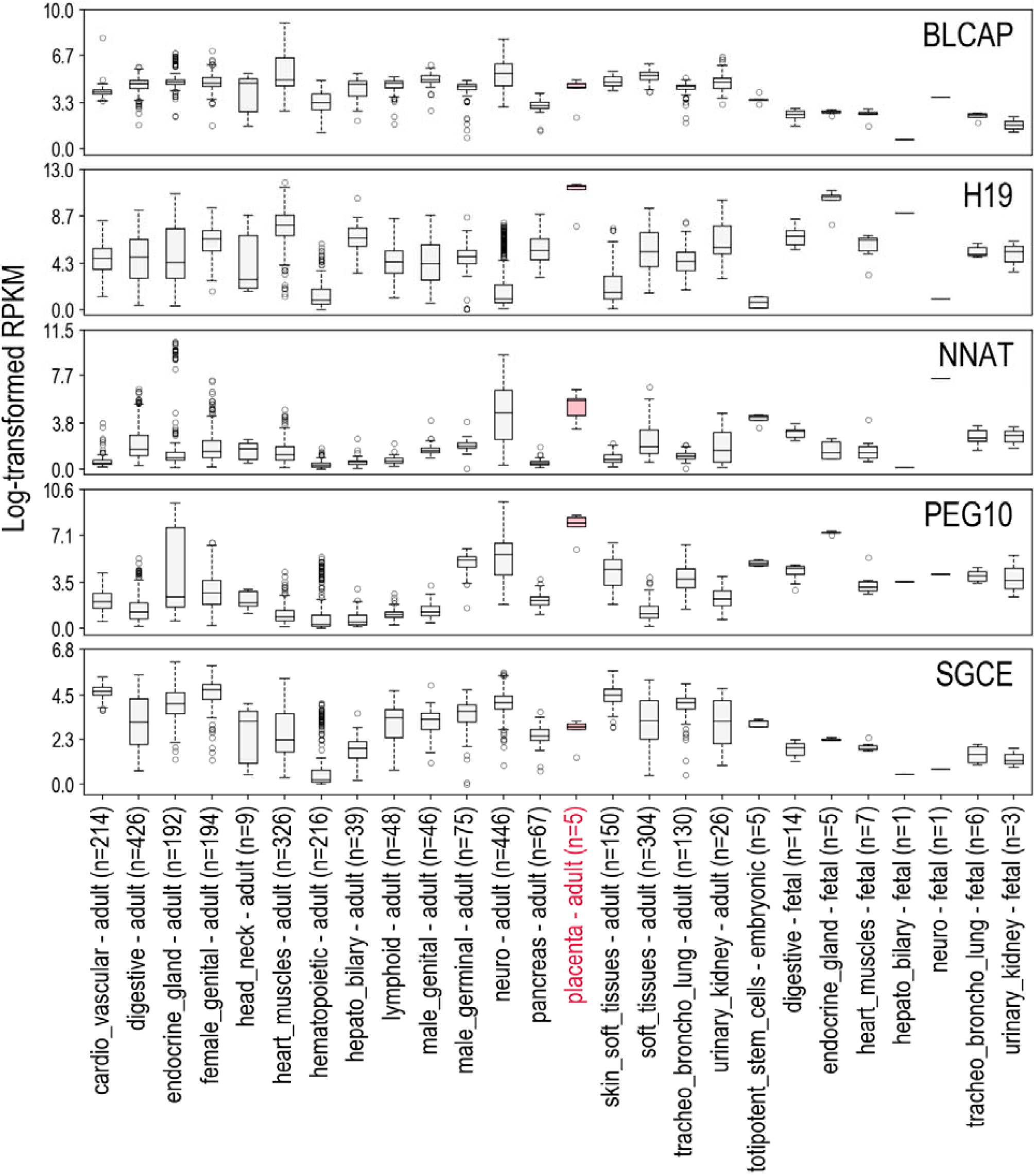
Expression of genes in normal human tissues from RNA-seq data provided by NCBI Sequence Read Archive (SRA) datasets PRJNA280600, PRJEB4337, PRJEB2445, PRJNA270632, NCBI GEO datasets GSE70741, GSE53096 and GTEx database. The expression levels are represented as distributions of log-transformed RPKM (Read per Kilobase Million) values after addition of a pseudo count of 1 (log2(1+RPKM)). The solid lines in the boxplots correspond to the median expression values. Expressions in placenta are marked in pink.

Table S1: Results from the Epigenome Wide Association Study (EWAS): 3,415 CpGs differentially methylated between the three groups of women (nonsmokers, current smokers or former smokers) (p-value corrected for False Discovery Rate (FDR) <0.05)

Table S2: Results from the regional analysis using comb-p: 344 Differentially Methylated Regions including 1741 CpGs

Table S3: Comparison of the number of CpGs found significantly associated with smoking between the present EDEN study and previous studies conducted on placenta and cord blood

Table S4: Metadata of analyzed ENCODE files.

## Acknowledgments

Most of the computations presented in this paper were performed using the CIMENT/GRICAD infrastructure (https://gricad.univ-grenoble-alpes.fr). We thank the participants of the EDEN cohort. We are grateful to the midwife research assistants (L. Douhaud. S. Bedel. B. Lortholary. S. Gabriel. M. Rogeon. and M. Malinbaum) for data collection and P. Lavoine for checking, coding, and entering data. The EDEN mother-child cohort study group includes: I Annesi-Maesano, JY Bernard, J Botton, M-A Charles, P Dargent-Molina, B de Lauzon-Guillain, P Ducimetière, M de Agostini, B Foliguet, A Forhan, X Fritel, A Germa, V Goua, R Hankard, B Heude, M Kaminski, B Larroque, N Lelong, J Lepeule, G Magnin, L Marchand, C Nabet, F Pierre, R Slama, MJ Saurel-Cubizolles, M Schweitzer, O Thiebaugeorges.

## Author contributions

JL, SK, SR, designed the study and wrote the manuscript. EBF, ES, FC, JL, MB performed the statistical and omics analyses. MAC, BH, VS, RS, JT, DV and the EDEN mother-child cohort study group provided guidance and interpretation of findings. All authors read, revised and approved the manuscript.

## Data and materials availability

The data are not publicly available due to them containing information that could compromise research participant privacy/consent.

## Web resources

NCBI Dataset: We used RNA-seq data provided by the NCBI Sequence Read Archive (SRA) datasets PRJNA280600, PRJEB4337, PRJEB2445, PRJNA270632 and NCBI GEO datasets GSE70741, GSE53096.

Genotype-Tissue Expression Project https://gtexportal.org/

Corresponding publication: https://www.ncbi.nlm.nih.gov/pmc/articles/PMC4010069/

We used RNA-seq data available on https://gtexportal.org/home/datasets

ENCODE Dataset: We used Chip-seq data provided by the ENCODE datasets https://www.encodeproject.org (Corresponding Publication: https://www.ncbi.nlm.nih.gov/pmc/articles/PMC3439153/)

### Transparency declaration

The lead author (the manuscript’s guarantor) affirms that this manuscript is an honest, accurate, and transparent account of the study being reported; that no important aspects of the study have been omitted; and that any discrepancies from the study as planned (and, if relevant, registered) have been explained.

## Patient and Public Involvement

Patients or the public were not involved in the design, or conduct, or reporting, or dissemination of our research.

The results are disseminated to study participants through newsletters and the study website (http://eden.vjf.inserm.fr).

## Supplementary Materials

**Supplementary tables S1 to S4 are provided in an excel file**

